# *Arabidopsis thaliana* Zn^2+^-efflux ATPases HMA2 and HMA4 are required for resistance to the necrotrophic fungus *Plectosphaerella cucumerina* BMM

**DOI:** 10.1101/2020.08.10.243014

**Authors:** Viviana Escudero, Álvaro Castro-León, Darío Ferreira Sánchez, Isidro Abreu, María Bernal, Ute Krämer, Daniel Grolimund, Manuel González-Guerrero, Lucía Jordá

## Abstract

- Zinc is an essential nutrient at low concentrations, but toxic at slightly higher ones. This could be used by plants to fight pathogens colonization.
- Elemental distribution in *Arabidopsis thaliana* leaves inoculated with the necrotrophic fungus *Plectosphaerella cucumerina* BMM (*PcBMM*) was determined and compared to mock-inoculated ones. Infection assays were carried out in wild type and long-distance zinc trafficking double mutant *hma2hma4*, defective in root-to-shoot zinc partitioning. Expression levels of genes involved in zinc homeostasis or in defence phytohormone-mediated pathways were determined.
- Zinc and manganese levels increased at the infection site. Zinc accumulation was absent in *hma2hma4. HMA2 and HMA4* transcription levels were upregulated upon *PcBMM* inoculation. Consistent with a role of these genes in plant immunity, *hma2hma4* mutants were more susceptible to *PcBMM* infection, phenotype rescued upon zinc supplementation. Transcript levels of genes involved in the salicylic acid, ethylene and jasmonate pathways were constitutively upregulated in *hma2hma4* plants.
- These data are consistent with a role of zinc in plant immunity not only of hyperaccumulator plants, but also of plants containing ordinary levels of zinc. This new layer of immunity seems to run in parallel to the already characterized defence pathways, and its removal has a direct effect on pathogen resistance.

## INTRODUCTION

Zinc concentration has to be kept within a very narrow range in all cells (Frausto da Silva & Williams, 2001; Outten & O’Halloran, 2001). Low zinc levels deprive the cell of the essential cofactor of around 10 % of its proteome (Andreini *et al.*, 2006; Broadley *et al.*, 2007), including enzymes involved in stress resistance and a large number of transcription factors. However, a slight excess of intracellular zinc results in toxicity, as zinc can interfere with the uptake of other essential transition metals or displace these in the active sites of enzymes (McDevitt *et al.*, 2011; Hassan *et al.*, 2017). This dual nature of zinc seems to be used by different organisms to fend-off invading microbes. Infected hosts may withhold growth-limiting nutrients from a pathogen to starve it and control its proliferation, in what has been known as nutritional immunity. For example, mammals remove zinc to combat bacterial and fungal infections (Kehl-Fie & Skaar, 2010; Grim *et al.*, 2020). Alternatively, host organisms may accumulate Zn either globally or locally in order to poison a pathogen. For example, zinc hyperaccumulator plants concentrate zinc to high levels in leaves, thus achieving some protection against herbivores, sap-feeding organisms and pathogenic microbes (Fones *et al.*, 2010; Kazemi-Dinan *et al.*, 2014; Stolpe *et al.*, 2017).

To date, there is only little direct evidence for zinc-mediated immunity in non-hyperaccumulator plants. For instance, *zur* (zinc uptake regulator) mutants in plant pathogens *Xanthomonas campestris* or *Xylella fastidiosa* are less virulent than wild type strains (Tang *et al.*, 2005; Navarrete & De La Fuente, 2015). This is highly suggestive of zinc playing a role in the host plant defense strategy. Zur proteins reduce zinc uptake under excess conditions and, in some species, activate the zinc detoxification machinery (Mikhaylina *et al.*, 2018). It can be hypothesized that the reduced virulence of *zur* strains is due to the lack of protection against high zinc levels in plants. However, we do not presently know whether plants accumulate zinc cations at infection sites and, if so, how this operates at the molecular level.

Plant zinc homeostasis has been thoroughly studied in the model *Arabidopsis thaliana* (Olsen & Palmgren, 2014). Uptake from the rhizosphere into the root symplasm is very likely mediated by ZIP (Zrt1-like, Irt1-like Proteins) transporters, such as AtZIP1, AtIRT3, AtZIP4, and/or AtZIP9 (Korshunova *et al.*, 1999; Lin *et al.*, 2009; Assunção *et al.*, 2010). Transporting zinc in the opposite direction, MTPs (Metal Tolerance Proteins) are involved in zinc efflux from the cytosol, either into cellular compartments (storage or zinc metalation) or out of the cell. For instance, AtMTP1 and AtMTP3 participate in the sequestration of zinc into vacuoles (Arrivault *et al.*, 2018; Desbrosses-Fonrouge *et al.*, 2005) while AtMTP2 is involved in zinc delivery into the endoplasmic reticulum (Hanikenne *et al.*, 2008; Sinclair *et al.*, 2018). Root-to-shoot transport of zinc in the xylem is largely mediated by P_1B_-ATPases HMA2 and HMA4 (Hussain *et al.*, 2004). These partially redundant Zn^2+^-ATPases are localized in the plasma membrane of vascular cells, where they would be transporting Zn^2+^ from the cell cytosol into the apoplast (Eren & Argüello, 2004; Hussain *et al.*, 2004). An *hma2hma4* double mutant has increased zinc levels in roots and lowered zinc levels in shoots, associated with complex alterations in transcript levels of zinc homeostasis genes (Sinclair *et al.*, 2018). Consistent with a role in zinc nutrition and distribution from roots to shoots, *hma2hma4* plants largely recover the wild type phenotype upon exogenous application of zinc (Hussain *et al.*, 2004). Transcription factors bZIP19 and bZIP23 control local transcriptional responses to zinc deficiency (Assunção *et al.*, 2010), but systemically regulated transcriptional zinc deficiency responses are independent (Sinclair et al., 2018). If zinc is an integral part of plant innate immunity, it would be expected that zinc homeostasis genes, particularly those involved in long-distance metal allocation, were upregulated upon pathogen invasion.

In this work, we use synchrotron-based X-ray fluorescence (S-XRF) to show that zinc levels are increased at the infection site in *A. thaliana* leaves 48 hours after being inoculated with the necrotrophic fungus *Plectosphaerella cucumerina* BMM (*PcBMM*). This increase in local zinc levels requires *HMA2* and *HMA4* as indicated by the lack of zinc accumulation in double *hma2hma4* mutants. The enhanced susceptibility of *hma2hma4* double mutant plants *PcBMM* further supports a role of zinc in plant immunity and resistance to the necrotrophic fungus *PcBMM*.

## MATERIALS AND METHODS

### Plant material and growth conditions

Wild type *A. thaliana* Columbia-0 (Col-0) ecotype was used in this study as well as the following lines in the Col-0 background: *hma2-4*, *hma4-2*, double mutant *hma2hma4* (Hussain *et al.*, 2004), *agb1-2* (Ullah *et al.*, 2003), and *irx1-6* (Hernandez-Blanco *et al.*, 2007). The *35S:HMA2* line was in *hma2hma4* background (Hussain *et al.*, 2004). Plants were sown in a mixture of peat:vermiculite (3:1), covered with sterilized sand and grown in growth chambers under short day conditions (10 hours light photoperiod and ~150μEm^−2^s^−1^) at 20-22°C. For infection experiments with *PcBMM*, plants were grown in growth chambers at 22-24°C. To perform Zn complementation assays, plants were watered with 1 mM zinc twice a week.

### Synchrotron-based X-ray Fluorescence assays

Leaves of 16-day-old *A. thaliana* were inoculated with a 2.5 μl drop of either sterilized water (mock) or with a suspension of 4 × 10^6^ *PcBMM* spores/ml. Subsequently, plants were maintained under a saturated atmosphere (80-85 % relative humidity) and short-day conditions. Samples were collected prior to inoculation (0 hours-post-inoculation, hpi) or at 48 hpi. Fifteen plants per time point, treatment, and genetic background were generated. At harvest, mock or *PcBMM*-inoculated leaves were immediately covered with ultralene membrane and flash-frozen in isopentane chilled with liquid nitrogen.

The elemental spatial distribution was characterized by synchrotron radiation scanning micro-XRF at the Swiss Light Source (SLS; microXAS beamline, Villigen, Switzerland) in cryo-conditions (liquid nitrogen cryojet). Data were acquired with a micro-focused pencil beam with a size of about 2 μm diametre. The excitation energies of 9.7 keV and 9.8 keV were used, which allowed the detection of elements between Si and Zn (K lines). XRF spectra were treated with PyMca software (Solé *et al.*, 2007). Incoming flux and transmitted intensities were also recorded using a micro ionization chamber and a silicon carbide diode, respectively, allowing to analyze absorption contrast simultaneously together with the XRF signal. The two-dimensional projection maps were recorded using 50 μm of step size and a dwell time of 400 ms per pixel. Two samples were analyzed through scanning XRF approach. The lateral step-size of 5 μm was used for the scanning tomography scans. 120 lateral projections equally spaced over 180° were measured for each of the 6 scans, each one at a different height of the sample. The tomography scan dataset was analyzed using home-made python codes, using the Astra Toolbox library (van Aarle *et al.*, 2015; van Aarle *et al.*, 2016), the SIRT method and parallel beam GPU code (Palenstijn *et al.*, 2011).

### Pathogenicity Assays

Resistance assays with *PcBMM* were performed as described (Escudero *et al.*, 2019). Briefly, *PcBMM* spores (4 × 10^6^ spores/ml) were sprayed onto 16-day-old *A. thaliana* leaves grown under short day conditions. A minimum of 20 plants per genotype were used in each experiment. *agb1-2* (Llorente *et al.*, 2005) and *irx1-6* (Hernández-Blanco *et al.*, 2007) plants were used as susceptible and resistant controls, respectively. Plants were kept at high relative humidity for the remaining duration of the experiment. At the indicated times, shoots from at least 4 plants were collected and gDNA extracted to determine relative *PcBMM* biomass by qPCR using specific primers for *β-tubulin* from *PcBMM* and *UBC21* (*At5g25760*) from *A. thaliana* to normalize (Table S1). These assays were repeated in triplicate.

### Gene expression experiments

Gene expression was determined in 48 hpi sprayed-infected and mock-inoculated 16-days-old *A. thaliana* plants. Shoots and roots were collected (from at least 8 plants per independent experiment) and RNA extraction, cDNA synthesis, and qRT-PCRs were performed as reported (Jordá *et al.*, 2016). Oligonucleotides used are listed on Table S1. Gene expression was normalized with the house-keeping gene *UBC21*. The Ct values of three independent experiments were used to calculate the gene expression using the 2^(ΔCt) method (Schmittgen & Livak, 2008).The results were represented as n-fold relativized with mock plants. These assays were carried out in triplicate.

### Statistical Tests

Data were analyzed by Student’s unpaired *t*-test to calculate statistical significance of observed differences. Test results with p-values < 0.05 were considered as statistically significant.

## RESULTS

### Enhanced zinc accumulation at leaf *PcBMM* infection site of wild-type plants is abolished in *hma2 hma4* double mutants

To determine whether high levels of zinc and other transition are altered upon infection, S-XRF studies were carried out in cryofixed *PcBMM*-infected leaves of wild-type plants (Col-0). In all samples, due to the abundance in cell walls, calcium distribution was used to define the general leaf shape as well as to indicate the position of *PcBMM* mycelium in the leaf, as Ca^2+^ influxes are one of the hallmark early events after pathogen perception. Indeed, at 48 hours post inoculation (hpi), a high-density calcium-rich spot could be observed in wild type *A. thaliana* leaves, but not in mock-inoculated ones (Fig. 1), coincident with the position in which the fungal hyphae were growing. Interestingly, marked increases in zinc and manganese concentrations were also detected in the same positions, although at lower magnitudes and with a pattern that might be associated to the leaf veins. Manganese and zinc-enrichment at the infection sites were also present at lower levels at the earlier 24 hpi time point (Fig. S1). Tomographic reconstructions of different fluorescence sections showed that both transition metals located to the surface of the leaf, where the spores germinated, and the mycelium was proliferating (Fig. S2). No other transition metal was observed at high levels at the time points analysed (Fig. S3).

**Figure 1.**
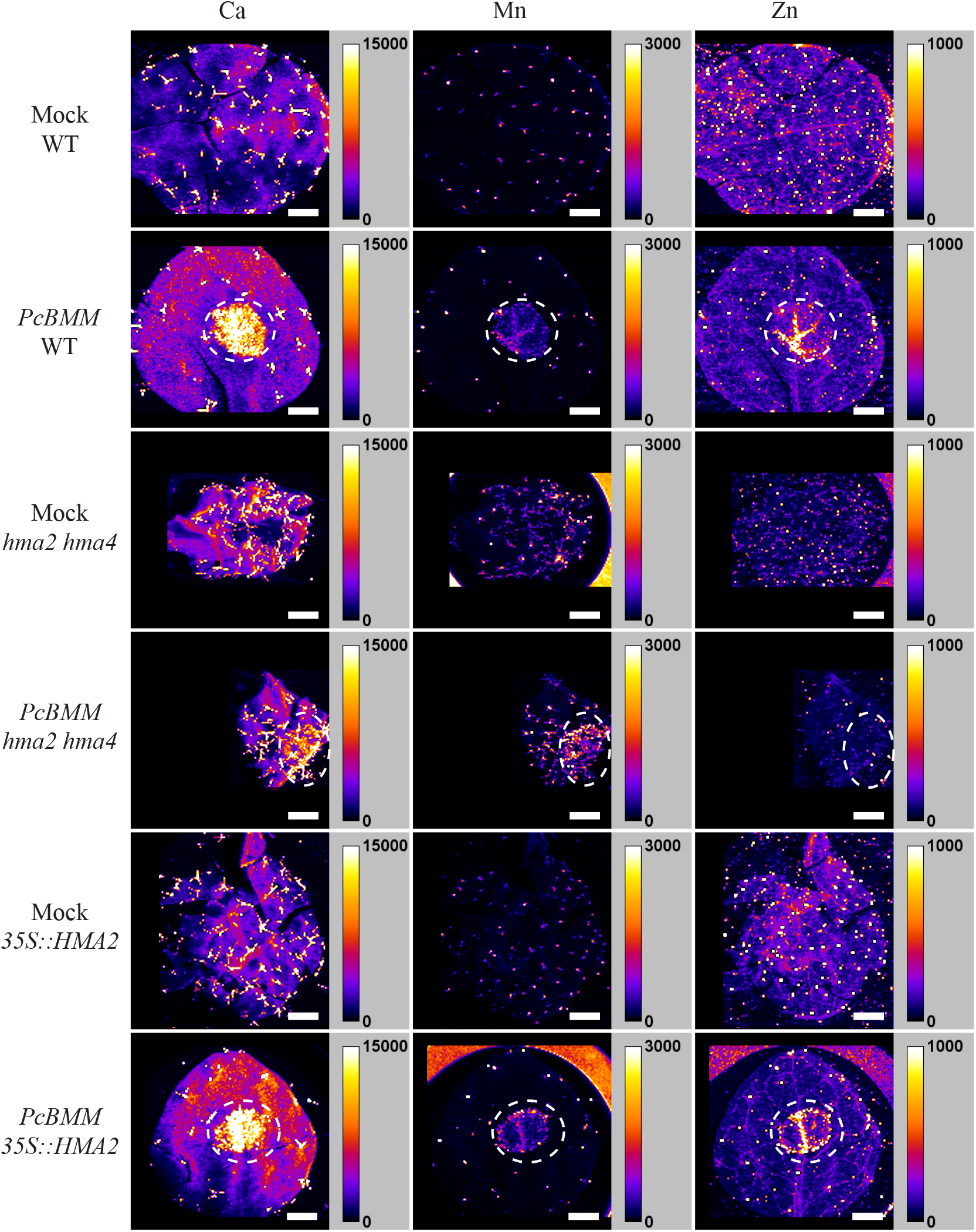
Zinc and manganese accumulate locally at the *PcBMM* infection site in *A. thaliana* leaves. Synchrotron-based X-ray fluorescence images of leaves of wild type Col-0 (WT), *hma2hma4* mutant, and the *hma2hma4* mutant expressing a wild type copy of the *HMA2* cDNA under a 35S promoter (*35S::HMA2*) 48 hours post inoculation with *PcBMM* or mock-treated. Left column shows the calcium distribution; centre, manganese; and right, zinc. Position of the calcium-rich spots is surrounded by the dashed line. Units indicate number of photon counts. Each image is the representative of three images taken from a randomly chosen leaf, each from a different Arabidopsis plant for each of the treatments and genotypes analysed.

The transporters HMA2 and HMA4 make the predominant contribution to root-to-shoot translocation of zinc (Hussain *et al.*, 2004). HMA2 also has a very modest Mn^2+^ transport capability (Eren & Argüello, 2004). Therefore, these two proteins are likely candidates of zinc, and perhaps manganese, accumulation at the infection site. Notably, 48 hpi *hma2hma4* mutant leaves did not show the localized enhanced zinc levels observed in Col-0 (Fig. 1), whereas accumulation of calcium and manganese were still abundant and reached similar levels to that of wild-type plants. Infection-induced zinc accumulation was restored to levels indistinguishable from the wild type in *hma2 hma4* plants into which *HMA2* had been reintroduced under the control of a 35S promoter.

### *HMA2* and *HMA4* are required for *A. thaliana* resistance to *PcBMM*

Enhanced zinc allocation to the infection site is suggestive of an up-regulation in the transcription levels of at least one of these Zn^2+^-ATPases. Real-time RT-PCR analyses of leaves at 48 hpi showed a consistent and significative induction of the transcription of both genes. *HMA2* transcript levels were up-regulated over two-fold in both shoots and roots of plants infected with *PcBMM* in comparison to mock-treated plants at 48 hpi (Fig. 2A). *HMA4* expression was highly up-regulated in roots of infected plants, but not in shoots (Fig. 2B). In contrast, transcript levels of other genes implicated in zinc transport and its regulation were not significantly changed in *PcBMM*-infected compared to mock inoculated plants (Fig. S4). This included genes with roles under zinc deficiency (*bZIP19*, *bZIP23*, *ZIP4*, *ZIP9*, or *MTP2*) as well as in detoxification (*MTP1*, *MTP3*) (Desbrosses-Fonrouge *et al.*, 2005; Arrivault *et al.*, 2006; Assunção *et al.*, 2010; Sinclair *et al.*, 2018).

**Figure 2.**
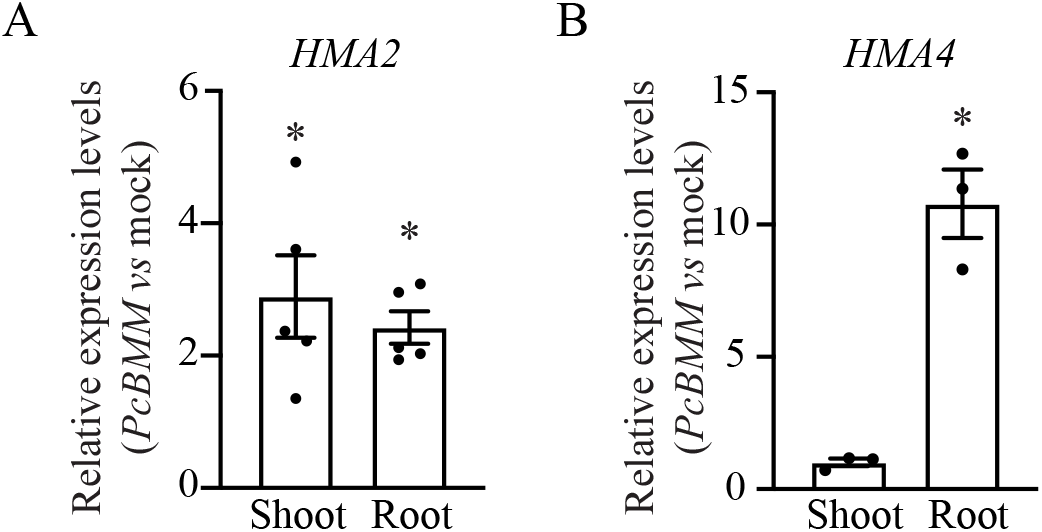
*HMA2* and *HMA4* are up-regulated upon *PcBMM* infection. (A) Expression of *HMA2* in 48 hpi shoots and roots normalized to mock inoculated plants. Data shows the mean ± SE of five independent infection assays, with tissues from 8-10 plants pooled per experiment. (B) Expression of *HMA4* in 48 hpi shoots and roots relativized to mock inoculated plants. Data shows the mean ± SE of three independent infection experiments, in each of them collecting 8-10 pooled plants. * indicates statistically significant difference from mock-infected plants according to Student’s *t*-test (*p*-value < 0.05).

These data are consistent with a role for *HMA2* and *HMA4* in zinc mobilization to the infection site, regulated at the transcript level, as part of the Arabidopsis innate immune response. To further test this possibility, wild type, *hma2*, *hma4*, *hma2hma4*, and *35S::HMA2* in *hma2hma4* background were sprayed-inoculated with *PcBMM*. Fungal biomass was determined in leaves at 5 days-post-inoculation (dpi) by the relative ratio of fungal *vs* plant gDNA using qPCR (Fig. 3A). As expected, *hma2hma4* had a higher pathogen proliferation, similar to what is observed in the hypersusceptible mutant *agb1-2.* Single mutants *hma2* and *hma4* had a very similar response to *PcBMM* than Col-0 wild-type plants, indicating that HMA2 and HMA4 have redundant functions in immune responses to this fungus. Of note, *35S::HMA2 hma2hma4* plants overexpressing *HMA2* recovered the disease resistance level of wild-type plants (Col-0), further confirming the functionality of *HMA2* in disease resistance to *PcBMM*. These observations were also supported by visual evaluation of the macroscopic symptoms in infected plants compared to the mock-inoculated controls (Fig. 3B).

**Figure 3.**
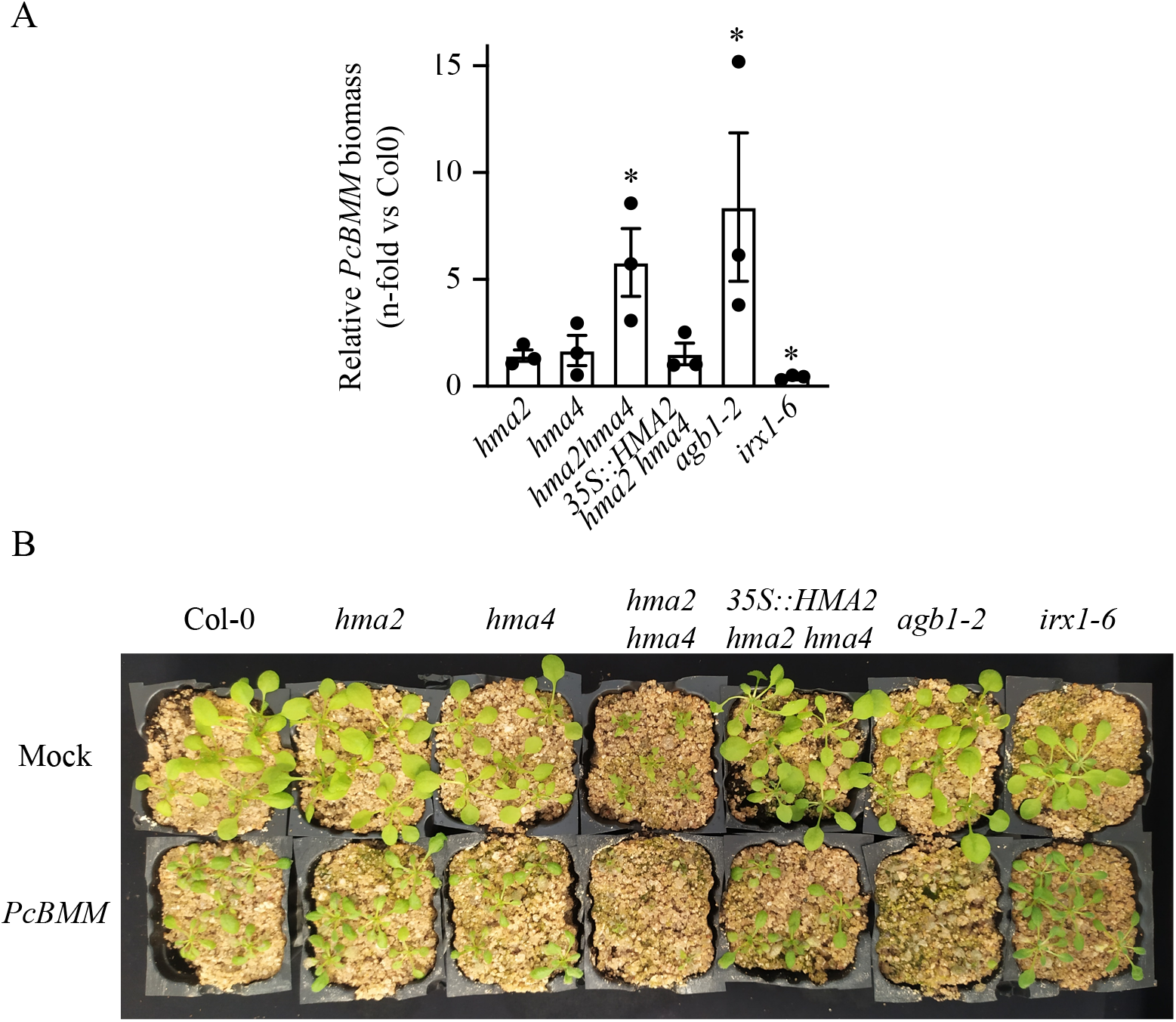
Mutants impaired in Zn^2+^-efflux ATPases HMA2 and HMA4 are more susceptible to infection by the necrotrophic fungus *PcBMM*. (A) quantification of *PcBMM* biomass by qPCR in the indicated genotypes at 5 dpi upon spray-inoculation with a suspension of 4×10^6^ spores/ml of the fungus. *agb1-2* and *irx1-6* plants were included as susceptible and resistant controls, respectively. Data shown are relative levels of fungal *β-tubulin* to Arabidopsis *UBC21*, normalized to the values of wild-type (Col-0) plants. Represented data are means ± SE, of three independent infection assays. Asterisks indicate statistically significant difference from the wild type according to Student’s *t*-test (*p*-value < 0.05). (B) Macroscopic symptoms of mock and *PcBMM*-inoculated plants at 8 dpi. Photographs are from one experiment representative of three independent experiments.

The growth defect of *hma2hma4* mutants has been demonstrated to be restored by supplementing the irrigation water with zinc (Hussain *et al.*, 2004). Similarly, when *hma2hma4* plants were watered with 1 mM zinc, their susceptibility to *PcBMM* was ameliorated. Double mutant plants watered with additional zinc had a severe reduction in fungal proliferation at 5 dpi (Fig. 4A), and their defence response was completely restored to wild type levels. Also, the overall look of these plants was much healthier than when no zinc was added to irrigation water (Fig. 4B).

**Figure 4.**
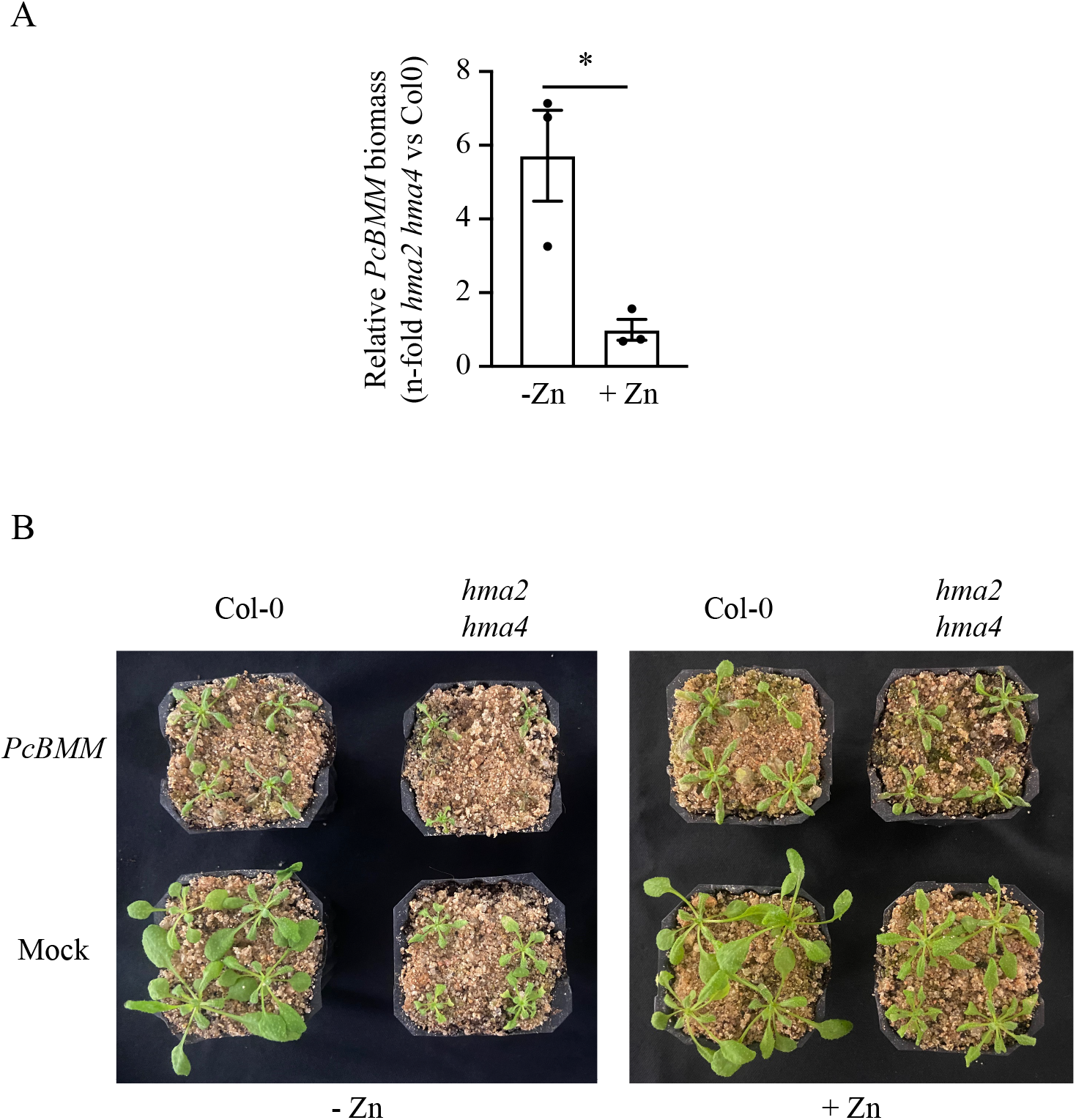
Application of exogenous zinc restores wild type infection levels in *hma2 hma4* plants. (A) quantification of *PcBMM* biomass by qPCR of the indicated genotypes at 5 dpi with a spray-inoculation with a suspension of 4×10^6^ spores/ml of the fungus. Data shown are relative levels of fungal *β-tubulin* to Arabidopsis *UBC21*, normalized to Col-0 values. -Zn indicates no added zinc in the watering solution and + Zn indicates 1 mM zinc sulphate used in the watering solution twice per week. Represented data are means ± SE, of three independent infection experiments. Asterisks indicate statistically significant difference from the wild type according to Student’s *t*-test (*p*-value < 0.05). (B) Macroscopic symptoms of mock and *PcBMM*-inoculated plants at 8 dpi. Experiments were performed three times with similar results. Photographs are from one experiment representative of three independent experiments.

### Mutations in *HMA2* and *HMA4* lead to transcriptional activation of defence genes

Plant hormones play central roles in modulating defence resistance mechanisms upon pathogen perception (Bürger & Chory, 2019). Among them, the ethylene (ET), jasmonic acid (JA), abscisic acid (ABA) and salycilic acid-mediated pathways orchestrate a complex network that contributes to plant immunity. Alterations in any of these phytohormone pathways diminishes resistance to pathogens, including the necrotrophic fungus *PcBMM* (Nawrath & Métraux, 1999; Adie *et al.*, 2007; Hernandez-Blanco *et al.*, 2007; Sanchez-Vallet *et al.*, 2010). To test whether the enhanced susceptibility of *hma2hma4* plants was due to alterations in any of the main defence signalling pathways, we quantified the expression of the following signature genes *PR1* (SA), *PDF1.2* (ET and JA), *LOX2* (JA) and *RD22* (ABA) in wild type and mutant plants under mock and *PcBMM*-inoculated conditions. Figure 5A shows that no significant differences could be observed in the expression levels of the *RD22* gene between *hma2hma4* and wild type plants, while, *PR1*, *PDF1.2* and *LOX2* were highly expressed in mock inoculated *hma2hma4* compared to Col-0 plants. This up-regulation in *hma2hma4* was maintained for *PR1* when the plants were infected with the necrotrophic fungus (Fig. 5B), but no additional induction was observed for *LOX2* or *RD22.* Besides, the *hma2hma4* mutant showed a strong repression in the expression of the *PDF1.2* gene after infection with the pathogen.

**Figure 5.**
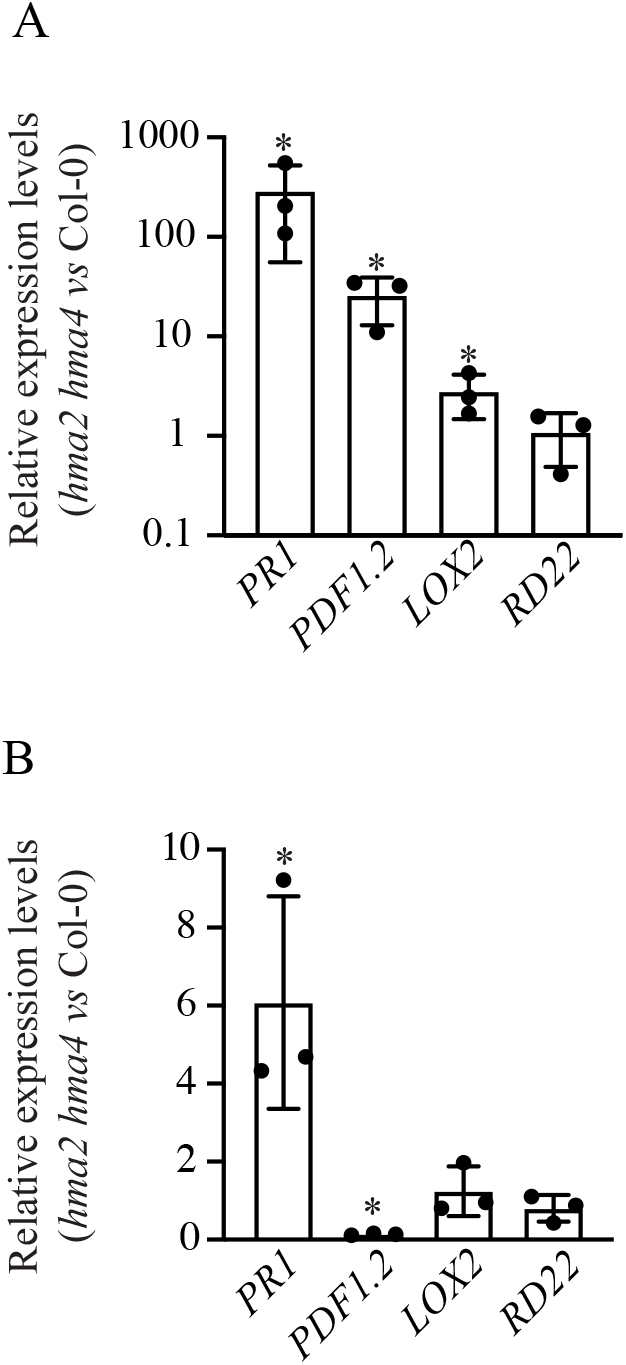
SA and JA/ET-signalling pathways are upregulated in *hma2 hma4* shoots. (A) Transcript levels of marker genes for the salicylic acid pathway (*PR1*), jasmonic acid pathway (*LOX2*), ethylene and jasmonate pathways (*PDF1.2*) and abscisic acid pathway (*RD22*) in shoots of mock-inoculated *hma2 hma4* relative to wild-type (Col-0) plants. Shown are mean ± SE of three independent infection experiments, with tissues pooled from 8-10 plants per experiment. (B) Expression analysis of marker genes for the SA, JA, JA/ET and ABA signalling pathways in shoots of *hma2 hma4* relative to wild-type (Col-0) plants at 48 hpi upon inoculation with a spore suspension of *PcBMM*. Shown are mean ± SE of three independent infection experiments, with tissues pooled from 8-10 plants per experiment.

## DISCUSSION

Life walks a narrow edge between zinc toxicity and zinc deficiency (Frausto da Silva & Williams, 2001). This can be used to combat invading microbes. Zinc deficiency can be produced locally to starve the invader (Kehl-Fie & Skaar, 2010), while it might also be increased to toxic levels to eliminate it (Fones *et al.*, 2010). Both strategies seem to be used in innate immunity. Zinc deficiency is favored in mammal immune systems (Kehl-Fie & Skaar, 2010; Hood & Skaar, 2012), while plants, and not only the zinc hyperaccumulators (Fones *et al.*, 2010; Kazemi-Dinan *et al.*, 2014; Stolpe *et al.*, 2017), seem to prefer the toxicity approach.

Our data shows that zinc and manganese are locally increased at *PcBMM* infection sites of leaves. The *hma2hma4* mutant is unable to mount a local increase in zinc levels at the infection site, and it is more susceptible to infection by the necrotrophic fungus *PcBMM*. Wild-type levels of resistance were restored in the *hma2hma4* mutant by application of exogenous zinc or constitutive overexpression of *HMA2* in the *hma2hma4* background. Two alternative and not incompatible explanations can be offered for these observations: i) Arabidopsis is using large amounts of zinc-proteins to combat *PcBMM* infection, or ii) Arabidopsis is using zinc to poison the invader. Within the first hypothesis, zinc limitation in the *hma2 hma4* mutants could reduce the activity of one or several zinc-proteins required for resistance to *PcBMM*. In this sense, PDF1.2 and other defensins have been shown to be able to bind zinc and to play a role in plant immunity and plant zinc tolerance (Shahzad *et al.*, 2013). However, this explanation would require the expression and concentration at the infection site of a large amount of zinc-proteins to account for the large increase of zinc at the infection site. Furthermore, it does not explain why pathogen strains in high risk of zinc toxicity are less virulent (Tang *et al.*, 2005; Navarrete & De La Fuente, 2015).

The alternative hypothesis suggests that a large portion of the zinc observed at the infection site would be free, hydrated. HMA2 and HMA4 would increase zinc concentrations to toxic levels for *PcBMM*. In this scenario it would be expected that the ability to detoxify this element would provide a competitive edge, what agrees with reports that plant pathogens require zinc detoxification systems for efficient virulence (Tang *et al.*, 2005; Navarrete & De La Fuente, 2015). More recently, it has been shown that a *PcBMM* CDF/MTP gene (PcBMM_CBGP_AIM006405) is induced when infecting Arabidopsis leaves compared to free-living conditions (Muñoz-Barrios *et al.*, 2020). Considering that CDF/MTP genes are involved in zinc detoxification, this is further indication of *PcBMM* facing high zinc levels at the infection site. The general pattern of pathogen protection against excess zinc as part of bacterial and fungal infection processes, indicates that zinc-mediated immunity would be a more general process not only limited to *PcBMM.* This use of zinc in plant immunity contrasts to what has been predominantly reported with animal pathogens, in which the ability to bind and uptake zinc with high affinity is a necessary requirement (Neumann *et al.*, 2017; Zackular *et al.*, 2020).

Manganese also accumulates at the infection site at similar time and at higher concentrations as zinc, what could indicate a role in *PcBMM* resistance. Further analyses in manganese transporter mutants might also yield similar results for manganese-mediated immunity. At the timepoints analysed in our S-XRF experiments, we did not observe any major changes in the distribution of iron or copper, in spite of existing literature indicating that it should be present. Upon phyopathogenic enterobacteria attack, Arabidopsis removes iron from the infection site to starve the invader (Aznar *et al.*, 2014; Aznar *et al.*, 2015), while copper levels should be increased as indicated by the loss of virulence in Arabidopsis of *Pseudomonas aeruginosa* that lose some of their copper-detoxification systems (González-Guerrero *et al.*, 2010). It is possible that infection-dependent changes in iron or copper localization in Arabidopsis leaves occurred, but were below our detection limits or occurred at time points other than 48 hpi.

Arabidopsis zinc-mediated immunity do not seem to be under the control of the known regulatory pathways of zinc homeostasis. Out of all zinc homeostasis genes tested, transcript levels responded to *PcBMM* infection only for *HMA2,* with mild increases in both roots and shoots, and *HMA4,* with a large increase confined to roots. Transcript levels of genes contributing to root zinc uptake from soil were not increased in response to infection (*IRT3, ZIP1, ZIP14, ZIP19*). Similarly, expression levels of genes encoding the transcription factors controlling locally regulated zinc deficiency responses (*bZIP19* and *bZIP23*) were unchanged. Gene expression of *MTP2* reflecting a systemically regulated zinc deficiency response was also unaltered in response to infection. Zinc detoxification, or vacuolar zinc sequestration (*MTP1* and *MTP3*), as a protecting mechanism against zinc toxicity was not transcriptionally increased, either. The transcriptional upregulation in roots of the Zn^2+^-ATPases when the pathogen is only applied in shoots illustrates that some systemic signaling occurs.

Our results indicate that zinc-mediated resistance is a fundamental mechanism in Arabidopsis innate immunity, as mutants impaired in the Zn^2+^-ATPases HMA2 and HMA4 are highly susceptible to *PcBMM*, despite presenting an upregulation of three of the main defence signalling pathways (SA, JA and ET). The higher expression levels of *PR1*, *PDF1.2*, and *LOX2* would reflect an attempt by the host plant to compensate for the lack of zinc-mediated immunity. However, this compensatory mechanism would not be sufficient to control fungal colonization of the double mutant plants. These data illustrate the relevant role of zinc to combat *PcBMM*. It should be noted that *hma2hma4* mutants are unable to activate *PDF1.2* marker gene expression after pathogen inoculation, in contrast to wild plants, suggesting a possible defect in the ET/JA signalling pathways, required for *PDF1.2* regulation. However, the expression levels of *PDF1.2* in the double mutant prior infection were higher than in Col-0 plants, suggesting that the defective up-regulation of *PDF1.2* only take place after pathogen reception. Future work will be directed to unveiling the connection of zinc-mediated immunity with the complex phytohormone-mediated defence signaling pathways.

Regardless of the specific mechanism, it seems that zinc transport via HMA2 and HMA4 is important for plant immunity, and that zinc itself might control fungal infection, as supported by the use of zinc-protective measures in plant pathogens (Tang *et al.*, 2005; Navarrete & De La Fuente, 2015; Muñoz-Barrios *et al.*, 2020). In addition to this process, yet-to-be-unveiled zinc-proteins might also be participating in *PcBMM* tolerance. Since zinc is a limiting nutrient (Alloway, 2008), it is intriguing why zinc has an important role in resistance to a pathogen instead of a more plentiful element. Perhaps the answer lies in its scarcity, to which most organisms are typically adapted so that they tend to accumulate it. It could also be that zinc toxicity takes advantage of the iron nutritional immunity in plants (Aznar *et al.*, 2014; Aznar *et al.*, 2015). Upon invading the host, a pathogen would up-regulate their iron uptake systems to ensure sufficient iron supply in the host environment. At the same time, this would make a pathogen more sensitive to zinc, since many iron transporters permeate other divalent metals as secondary substrates (Guerinot, 2000; Forbes & Gros, 2001; Nevo & Nelson, 2006), particularly if present at sufficiently high concentrations. This model could also explain manganese accumulation at the infection site. Zinc-mediated immunity may open up new strategies against plant pathogens using proper application of zinc enriched fertilizers.

## ACKNOWLEDGEMENTS

This work has been financially supported by the “Severo Ochoa Programme for Centres of Excellence in R&D” from the Agencia Estatal de Investigación of Spain (grant SEV-2016-0672 (2017-2021) to the CBGP). In the frame of this program Viviana Escudero was hired with a postdoctoral contract. Álvaro Castro-León was supported by an Industrial Doctorate in partnership with Genomics4All awarded by Comunidad de Madrid (IND2019/BIO-17117). Isidro Abreu was the recipient of a Juan de la Cierva-Formación postdoctoral fellowship from Ministerio de Ciencia, Innovación y Universidades (FJCI-2017-33222). We acknowledge the Paul Scherrer Institut, Villigen, Switzerland for provision of synchrotron radiation beamtime at beamline microXAS, accessed gained through proposals 20190571 and 20180921. The research leading to part of the SXRF data has received funding from the European Union’s Horizon 2020 research and innovation programme under grant agreement number 730872, project CALIPSOplus. We would also like to acknowledge Dr. Antonio Molina for critical reading of the manuscript, and the rest of members of M. González-Guerrero laboratory at Centro de Biotecnología y Genómica de Plantas (UPM-INIA) for their support and feedback in preparing this manuscript.

## AUTHOR CONTRIBUTION

VE carried out most of the experimental work with AC-L assisting in some of the infection assays. DF, IA, and DG helped in the beamtime experiments, facilitating the set up and data analyses. MB and UK provided the zinc mutants used, participated in the experimental design of the zinc resistence assays, and edited the manuscript. VE, MG-G, and LJ conceived the project with input from MB, and UK. MG-G and LJ coordinated the work and wrote the manuscript with the figures being prepared by VE, except Fig. 1 and 2 being produced by DF. All authors have read and approved this manuscript.

**The following Supporting Information is available for this article:**

**Fig. S1** Localized zinc and manganese accumulation can be detected at the inoculation site at 24 hpi with *PcBMM* in *A. thaliana* leaves.

**Fig. S2** Zinc and manganese accumulation takes place in the epidermal cell layer.

**Fig. S3** Arabidopsis leaves do not accumulate iron or copper at the infection site with *PcBMM* at 48hpi.

**Fig. S4.** Expression levels of other zinc homeostasis genes in roots and shoots of mock-inoculated and *PcBMM*-infected plants.

**Table S1.** Primers used in this study.

