## Supplemental for "*Arabidopsis thaliana* Zn^2+^-efflux ATPases HMA2 and HMA4 are required for resistance to the necrotrophic fungus *Plectosphaerella cucumerina* BMM"

**SUPPLEMENTAL FIGURE S1**

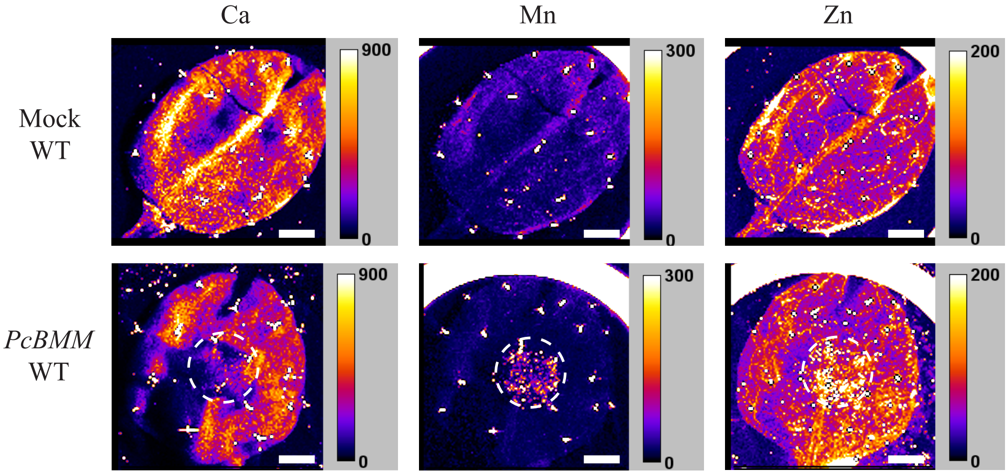


**Figure S1.** Localized zinc and manganese accumulation can be detected at the inoculation site at 24 hpi with *PcBMM* in *A. thaliana* leaves. Synchrotron-based X-ray fluorescence images of wild type Col-0 (WT) 24 hpi with *PcBMM* or mock-treated. Left column shows the calcium distribution; centre, manganese; and right, zinc. Position of the manganese-rich spots spots is surrounded by the dashed line. Units indicate number of photon counts. Scale bars = 1 mm

### SUPPLEMENTAL FIGURE S2

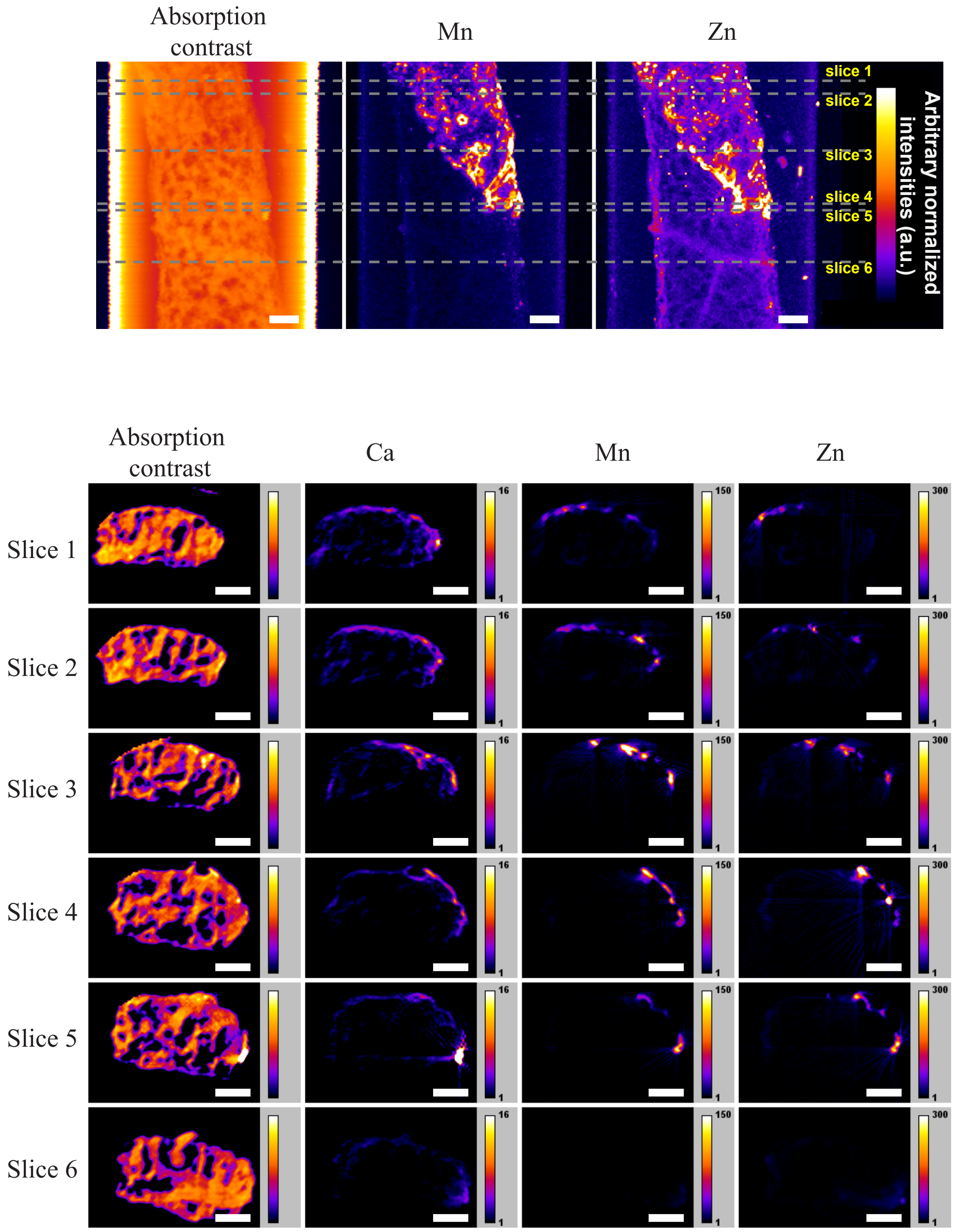


**Figure S2.** Zinc and manganese accumulation takes place at the epidermal cells. Top panel, tomographic reconstruction of synchrotron-based X-ray fluorescence images of wild type Col-0 48 hpi with *PcBMM*. Left image shows the absorption contrast image; centre, manganese; and right, zinc. Individual slices are shown in the lower panel. Left-most image is the absorption contrast image, next to it is calcium distribution, followed by manganese signal and zinc image at the right-most column. Scale bars = 100 µm.

### SUPPLEMENTAL FIGURE S3

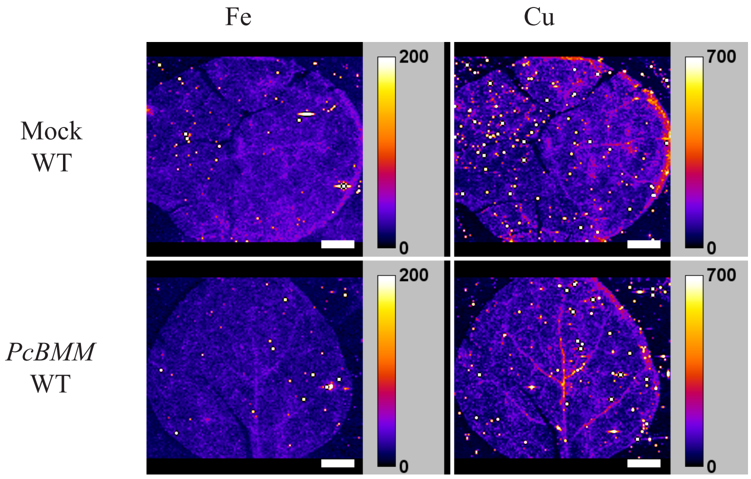


**Figure S3.** Arabidopsis leaves do not accumulate iron or copper at the infection site with *PcBMM* at 48hpi. Synchrotron-based X-ray fluorescence images of wild type Col-0 (WT) 24 hpi with *PcBMM* or mock-treated. Left column shows the iron distribution; and right, copper. Units indicate number of photon counts. Scale bars = 1 mm.

**SUPPLEMENTAL FIGURE S4**

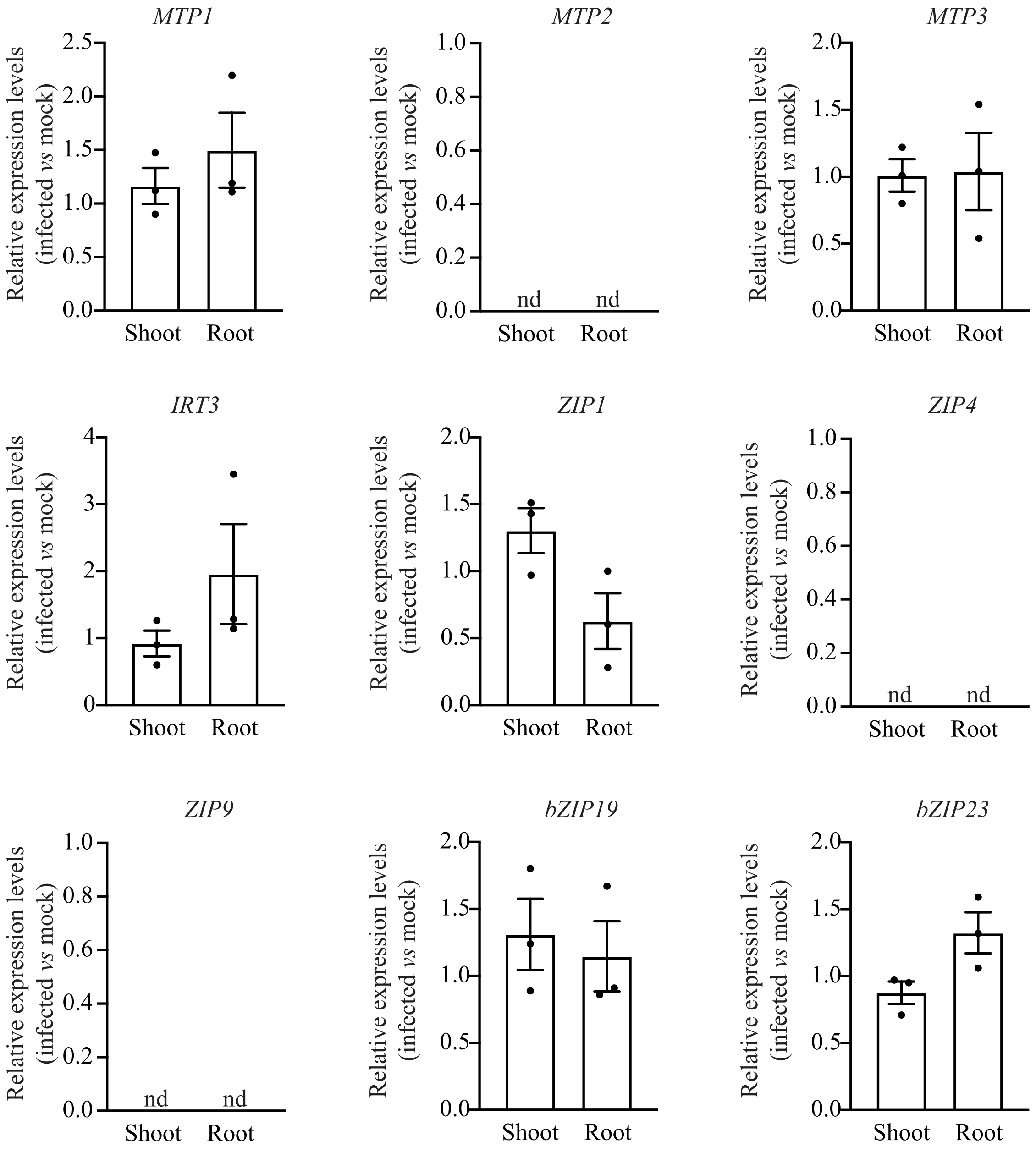


**Figure S4.** Expression levels of other zinc homeostasis genes in roots and shoots of mock-inoculated and *PcBMM*-infected plants. Expression of the indicated genes in 48 dpi shoots and roots relativized to mock inoculated plants. Data shows the mean ± SE of three independent infection assays, in each of them collecting 4 pooled plants.

**Table S1.** Primers used in this study.

| Gene | identity | Oligos 5´- 3´ | Oligos 3´- 5´ |
|---|---|---|---|
| *β-Tub 1* | Pc*BMM* | CAAGTATGTTCCCCGAGCCGT | GAAGAGCTGACCGAAGGGACC |
| *bZIP19* | AT4G35040 | ACGTGCTTCCATGTCCACACC | AACCGCTTCCCGGTTTCCCA |
| *bZIP23* | AT2G16770 | AGCGGCTTCGTTGGAGGATGA | AGGCACGTTTGTAACCGCAGG |
| *HMA2* | AT4G30110 | GGTTAGGAGTACAAAATATAAGTGAG | TTGTAGGAGCTACGCTAAAGA |
| *HMA4* | AT2G19110 | AGAGAGCACGAATTGTTCCACG | GCCTGATACCACCAAGCTAGCA |
| *IRT3* | AT1G60960 | TCCGGCTGAGAGCGAGTCCA | TTTCGCCCTCACAACCCCGA |
| *LOX2* | AT3G45140 | ATCAACAAGCCCCAATGGAA | CGGCGTCATGAGAGATAGCAT |
| *MTP1* | AT2G46800 | TGTATCGCCGTCGTGCTGTGT | AGGAGTCGCTTCCCAGCCAG |
| *MTP2* | AT3G61940 | GCAAATCCGCGACAGAGTTATGGG | TGTCCGTGATCATGTCCAAGCACA |
| *MTP3* | AT3G58810 | ACGGTAGTGGTGAAGTGGAGGGA | TGATGATGCTCGGTTGCGGCT |
| *PDF1.2* | AT5G44420 | TTCTCTTTGCTGCTTTCGACG | GCATGCATTACTGTTTCCGCA |
| *PR1* | AT2G14610 | CGTCTTTGTAGCTCTTGTAGGTGC | TGCCTGGTTGTGAACCCTTAG |
| *RD22* | AT5G25610 | CTGTTTCCACTGAGGTGGCTAAG | TGGCAGTAGAACACCGCGA |
| *UBC 21 (cDNA)* | AT5G25760 | GCTCTTATCAAAGGACCTTCGG | CGAACTTGAGGAGGTTGCAAAG |
| *UBC 21 (gDNA)* | AT5G25760 | AAAGGACCTTCGGAGACTCCTTACG | GGTCAAGAATCGAACTTGAGGAGGTT |
| *ZIP1* | AT3G12750 | CTGTCGGCGATGGGGACTCTTA | CCGTGACTAGCGTGCGTGTGAA |
| *ZIP4* | AT1G10970 | GTCTCCATGCACGATCAGGCCT | TCAGTGACCAAAGCTCCAGGGC |
| *ZIP9* | AT4G33020 | TGGAGGGGCGTTGCATATCGTG | TGCCTCACACCACTGTCCAAGC |
